# Old Dog, New Tricks: Influenza A Virus NS1 and *In Vitro* Fibrillogenesis

**DOI:** 10.1101/2020.08.17.254060

**Authors:** A.A. Shaldzhyan, Y.A. Zabrodskaya, I.L. Baranovskaya, M.V. Sergeeva, A.N. Gorshkov, I.I. Savin, S.M. Shishlyannikov, E.S. Ramsay, A.V. Protasov, A.P. Kukhareva, V.V. Egorov

**Author notes:** Corresponding author: Dr. Yana A. Zabrodskaya.

## Abstract

The influenza NS1 protein is involved in suppression of the host immune response. Recently, there is growing evidence that prion-like protein aggregation plays an important role in cellular signaling and immune responses. In this work, we obtained a recombinant, influenza A NS1 protein and showed that it is able to form amyloid-like fibrils *in vitro*. Using proteolysis and subsequent mass spectrometry, we showed that regions resistant to protease hydrolysis highly differ between the native NS1 form (NS1-N) and fibrillar form (NS1-F); this indicates that significant structural changes occur during fibril formation. The discovery of the ability of NS1 to form amyloid-like fibrils may be relevant to uncovering relationships between influenza A infection and modulation of the immune response.

## INTRODUCTION

Influenza A nonstructural protein 1 (NS1) is a multi-functional protein, with a mass of 26 kDa, which is mainly involved in suppression host immune responses. Currently, a number of mechanisms are known that involve NS1 and which prevent the induction of an interferon response [1]. Despite an important role in influenza pathogenesis, many of the mechanisms by which NS1 affects cellular factors are unclear. Many viral proteins are prone to amyloid-like fibril formation [2], and some them have been shown to play roles in host cell interactions [3]. For the influenza A virus, functional amyloidogenicity has been shown for the PB1-F2 [4] protein, and an ability of PB1’s N-terminus to form fibrils has been shown as well [5]. It has recently been demonstrated that the NEP protein, which is a product of NS1 mRNA alternative splicing, is also capable of forming amyloid-like fibrils [6]. Fibrillogenesis is a phenomenon known to involve cell signaling in non-infected cells [7,8]. Studies have linked some viral proteins’ ability to undergo amyloid-like fibrillogenesis with such signaling. Amyloidogenic proteins display an ability to recruitment host proteins featuring similar primary structure [9] or to induce conformational changes in such host proteins [10]. This allows viral amyloidogenic proteins to modulate host protein function by changing their conformation or activity [11,12]. It should be noted that presence of full-length influenza A NS1 protein affects the CD8^+^ cell mediated immune response [13]. The effect may be induction of an immune response with an NS1 N-terminal fragment, or suppression of an immune response with a full-length protein. The role of the NS1 C-terminus in immunosuppression has been discussed [14]. In this study, we show that a recombinant NS1 protein is capable of forming amyloid-like fibrils *in vitro*.

## MATERIALS AND METHODS

### Genetic vector construction

Viral NS gene DNA sequence was obtained by PCR reaction, using the previously-constructed pHW2006 plasmid (encoding the NS segment of A/Puerto Rico/8/1934 H1N1 virus) as DNA template. The following primers were used: forward (5′-AACCTTCATATggATCCAAACACTgTg -3′), containing an Nde1 restriction site; and reverse (5′-TTAACTCgAgAACTTCTgACCTAATTgTTC -3′), containing an Xho1 restriction site. PCR was performed using a C1000 Thermal Cycler (Bio-Rad), Q5 High-Fidelity DNA Polymerase (NEB, M0491S), and 5x Q5 Reaction Buffer (NEB, B9027S) according to manufacturer protocols (53°C annealing temp.). Following electrophoretic separation in 1% agarose (in TBE), DNA bands were excised with subsequent purification of PCR products using a Cleanup Standard Kit (Evrogen, BC022). Restriction enzymes, namely ‘FastDigest XhoI’ and ‘FastDigest NdeI’ (Thermo Fisher Scientific, FD0695 and FD0585, respectively), were used to prepare sticky ends in insert and vector. The pET-22b+ vector (Novagen, USA) was used to make our construct (pET-NS1). Next, ligation was performed using T4 DNA Ligase (NEB, M0202L), and DH5a bacterial cells were electroporated with ligation mixes. Subsequent growth on ampicillin plates (100 ug/ml) enabled clone selection. After selection, pET-NS1 was purified using a Plasmid Miniprep Kit (Evrogen, BC021). Commercial plasmid sequencing was performed using T7 primers at Evrogen (Moscow, http://evrogen.com). Amino acid sequence corresponding to the coding region of the constructed plasmid is shown in Table 1. After sequence verification, the BL21(DE3) *E. coli* strain was transfected with pET-NS1 plasmid using standard protocols.

**Table 1.**
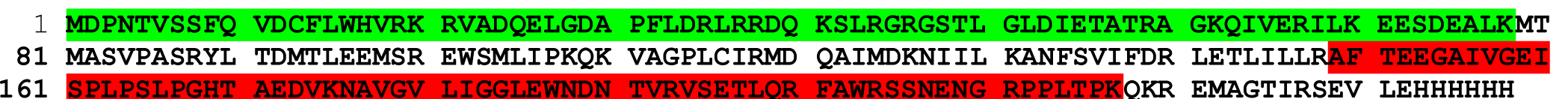
NS1-6xHis recombinant protein amino acid sequence. Green – N-terminal region, resistant to trypsin digestion in native NS1. Red – C-terminal region, resistant to trypsin digestion in fibrillar NS1.

### Protein expression

Three types of culture were used (pre-seed, overnight-seed, and production). Initial *E. coli* seed culture was incubated for 8 hr (200 rpm agitation, 37°C). Next, 200 µl of seed culture was transferred into 50 ml of LB medium (in 250-ml shaking flasks). The Luria– Bertani (LB) medium used for seed cultures contained (per 1 liter): 5 g yeast extract, 10 g tryptone, and 10 g NaCl. Ampicillin (100 mg/L) was used for growth of *E. coli* BL21(DE3) cells containing pET-NS1 to inhibit the growth of non-transformed cells. Growth under optimized conditions was carried out using a 10 L fermentor with agitation and airflow (BioFlo/CelliGen 115, New Brunswick). Overnight seed culture (100 ml) was used to inoculate 8 L of production medium (pH 7.4) containing: 15 g/L casein hydrolysate; 7.5 g/L yeast extract; 7 g/L NaCl; 5 g/L glucose; 1 g/L NH_4_Cl; 0.24 g/L MgSO_4_; 3 g/L KH_2_PO_4_X1H_2_O; 14.3 g/L Na_2_HPO_4_X2H_2_O; and 100 mg/L ampicillin. Following the appearance of several indicators (increase in dissolved oxygen and drop in pH), conditions were adjusted, and new nutrients were added containing non-glucose carbon sources: the temperature was manually decreased to 32°C; IPTG was added (1 mM final); additional carbon sources were added (casein hydrolysate 1.5 g/L, yeast extract 0.75 g/L, and glycerol 6.0 g/L); and incubation was continued. Culture pH was continuously maintained in a range from 6.9 - 7.0 by addition of phosphoric acid solution. Fermentation was continued until an optical density of 0.7-0.8 (at 600 nm) had been reached. Samples were taken every 1 hr (for 8 hrs), and cells were harvested using centrifugation at 4,500 *g* for 30 min (Eppendorf 5810r centrifuge). Cell pellets were stored at −20°C.

### Protein purification

Chromatographic purification of the NS1 protein was performed via immobilized metal affinity chromatography (IMAC), followed by buffer exchange via gel filtration on an AKTA Start chromatographic system (GE Healthcare, USA). Two grams of cell pellet were thawed in an ice bath and resuspended in sonication buffer (30 mM sodium phosphate buffer, pH 7.5, with 750 mM NaCl, 20 mM imidazole, 1 mM PMSF) using a volumetric ratio of 10 ml per 1 g of wet cells. Cells were lysed by probe sonication (MSE sonicator) with 10 cycles (each cycle: 1 min on, 1 min off). Cell debris was removed via centrifugation at 13,000*g* for 1 hour at +4°C (Eppendorf 5810r centrifuge). A 1 ml HisTrap FF Crude chromatography column (GE Healthcare, USA) was equilibrated with 5 ml of buffer (30 mM sodium phosphate buffer, pH 7.5, with 750 mM NaCl, 20 mM imidazole, 1 mM PMSF) at 1 ml/min. Supernatant obtained in the preceding step was loaded onto the column at the same flow rate. The column was washed with 20 ml of buffer (30 mM sodium phosphate buffer, pH 7.5, with 750 mM NaCl, 20 mM imidazole, 1 mM PMSF) at 1 ml/min. The bound protein was released with elution buffer (10 ml of 30 mM sodium phosphate buffer, pH 7.5, with 750 mM NaCl, 500 mM imidazole) at 1 ml/min., and 0.5 ml fractions were collected during elution.

Fractions containing target protein were identified via SDS-PAGE under reducing conditions, pooled, and filtered through a Sartorius syringe filter (33 mm diameter, PES membrane, pore size 0.22 μm). Recombinant NS1-6xHis protein was further purified by gel filtration using an AKTA pure chromatograph (GE Healthcare, USA). An XK 16/60 Chromatographic Column, packed with 120 ml of Sephadex S-200 HR sorbent (globular protein fractionation range 5 – 250 kDa), was washed sequentially with 120 ml of distilled water and 240 ml of 2x PBS.

Next, the recombinant protein was introduced into the column using a 5 ml injection loop and eluted with 180 ml of 2x PBS. During elution, 2.5 ml fractions were collected. Protein presence in eluate was monitored by absorption at a wavelength of 280 nm using a flow-through UV detector with a cell optical path length of 2 mm. Fractions were analyzed using SDS-PAGE under reducing conditions. Fractions containing pure target protein without detectable impurities were pooled. Column calibration was performed using a mixture of standard proteins (GE Gel Filtration Calibration Kit) according to manufacturer’s instructions.

### Polyacrylamide gel electrophoresis (PAGE)

Purified protein was analyzed via SDS-PAGE under reducing conditions using an AnykDa precast gel (Bio-Rad, USA) in a Mini Protean Tetra Cell device (Bio-Rad, USA), as described [15]. Following PAGE, the polyacrylamide gel was stained as follows: fixation for 15 minutes (10% acetic acid in 35% ethanol); then staining for 30 minutes (0.25% w/v Coumassie Brilliant Blue R250 (Bio-Rad, USA) in a 10% acetic acid, 25% ethanol solution). The gel was then incubated in destaining solution (10% acetic acid, 25% ethanol), until the background became transparent, and then transferred into water. Images were obtained using a ChemiDoc XRS+ digital image station (Bio-Rad, USA).

### Mass spectrometry

Enzymatic trypsin hydrolysis of proteins in gel was carried out without disulfide bond reduction and without iodoacetamide modification as described [16]. Briefly, excised gel fragments were washed twice for 15 minutes with dye removal solution (30 mM NH_4_HCO_3_ in 40% acetonitrile) and dehydrated with 100% acetonitrile. After complete evaporation of acetonitrile, 20 μg/ml porcine trypsin (Promega, USA) in 50 mM NH_4_HCO_3_ was added to the gel fragments. Enzymatic hydrolysis was carried out for 8 hours at 37°C. The reaction was stopped with a solution of 0.5% trifluoroacetic acid with 10% acetonitrile. The resulting tryptic peptides were mixed with an HCCA matrix (Bruker, Germany), applied to a GroundSteel target (Bruker, Germany), and analyzed on an ultrafleXtreme MALDI-TOF/TOF mass spectrometer (Bruker, Germany) in positive ion detection mode. For each spectrum, 5000 laser pulses were collected.

Recombinant protein sequence was added to local database as “NS1-6xHis”. Protein identification was performed using MASCOT (www.matrixscience.com) with simultaneous access to the SwissProt database (www.uniprot.org) and local database. As variable modifications, oxidation (M) and deamidation (NQ) were indicated. The mass error was limited to 20 ppm. Identification was considered reliable if the spectrum Score exceeded the threshold value (p < 0.05). In the Results section, the Mascot Score in comparison with threshold value is indicated using a slash. For example, ‘Score 100/70’ indicates: Mascot Score value 100, with threshold 70.

### Production of fibrils

NS1 fibrils were obtained as follows: one milliliter of NS1 recombinant protein (3 mg/ml in 2x PBS) was placed into a 1.5 ml Eppendorf tube (Eppendorf, Germany); and the tube mixed for 66 hours (55°C, 600 rpm) using a ThermoMixer Comfort orbital shaker (Eppendorf).

### Thioflavin T (ThT)

Thioflavin T (Sigma Aldrich, USA) stock solution (1 mM in PBS) was diluted to a 25 μM concentration in PBS immediately before measurements. To a 570 μl ThT solution, 30 μl of test sample was added, mixed thoroughly, and incubated for 15 minutes in darkness. For use as a negative control sample, the same volume of PBS buffer was added instead of the test sample. Thioflavin T fluorescence spectra were recorded on an Avantes AvaSpec 2048 instrument (Avantes, The Netherlands) with an LED radiation source having a maximum emission spectrum of 450 nm. Disposable plastic cuvettes (10×10 mm, Sarstedt, Germany) were used. Processing and visualization of spectra were performed using the Origin2015 software package.

### Congo red (CR)

Congo red (Sigma Aldrich, USA) was purified before use following previously described methods [17]. Two components were prepared: 50 μL of 500 μM Congo red solution in PBS; and 45 μL of NS1-6xHis solution/suspension (3 mg/ml, 110 μM). The components were mixed, and the final volume brought to 500 μL with PBS. The optimal CR:protein ratio was determined to be 5:1. Absorption spectra were recorded using an AvaSpec 2048 instrument (Avantes, The Netherlands) in optical cuvettes (Sarstedt, Germany) with a path length of 1 mm. PBS was used as a blank, and 50 μM CR solution in PBS was used for CR absorbance spectra measurement. To determine the presence of amyloid-like aggregates, a spectral difference approach was used. Specifically, such aggregates were considered present if subtraction of spectra (spectrum_(dye+protein)_ – spectrum_(dye)_) yielded a 540 nm peak [18]. Processing and visualization of spectra were performed using the Origin2015 software package.

### Electron microscopy

Ten microliters of sample was applied to a copper grid previously coated with collodion. After 1 minute of incubation, the sample was removed, and the grid was washed twice with water. For negative contrast, the grid with sample was incubated for one minute with 20 μl of sodium phosphotungstate. Excess acid was removed with a piece of filter paper. Electron micrographs were taken with a JEM 1011 electron microscope (JEOL Ltd., Japan) equipped with a Morada digital camera.

### Protein treatment with trypsin

In order to compare the protease accessibility of NS1-6xHis in native (NS1-N) and fibrillar (NS1-F) form, enzymatic hydrolysis with trypsin was performed in protein solution or fibrillar suspension, respectively. NS1-N or NS1-F (3 mg/ml) was mixed with trypsin solution (in 100 mM NH_4_HCO_3_) in equal volumes (NS1, trypsin) to reach trypsin:protein ratios of 1:20 or 1:100. The samples, including negative controls (mock-treated protein solutions), were incubated for 24 hours at 37°C. To stop trypsin hydrolysis reactions, 100 mM PMSF in ethanol was added up to a 1 mM final concentration. The samples were studied with SDS-PAGE (10-20% gradient gel) and mass spectrometry, as described earlier.

### Protein structure visualization

For visualization of NS1 structure, PyMol software was used [19]. The PDB access code is 4OPH [20,21].

## RESULTS AND DISCUSSION

The pET-NS1 plasmid was obtained as described in the Materials and Method section, and the insert sequence was verified using DNA sequencing. Protein expression and purification was performed as described. After gel-filtration (GF) (Figure 1a), fractions were analyzed using electrophoretic separation, and pure protein-containing fractions were combined (Figure 1a, inset). The amino acid sequence of the obtained NS1-6xHis protein was confirmed using mass spectrometry (Score 259/70, sequence coverage 95%).

**Figure 1.**
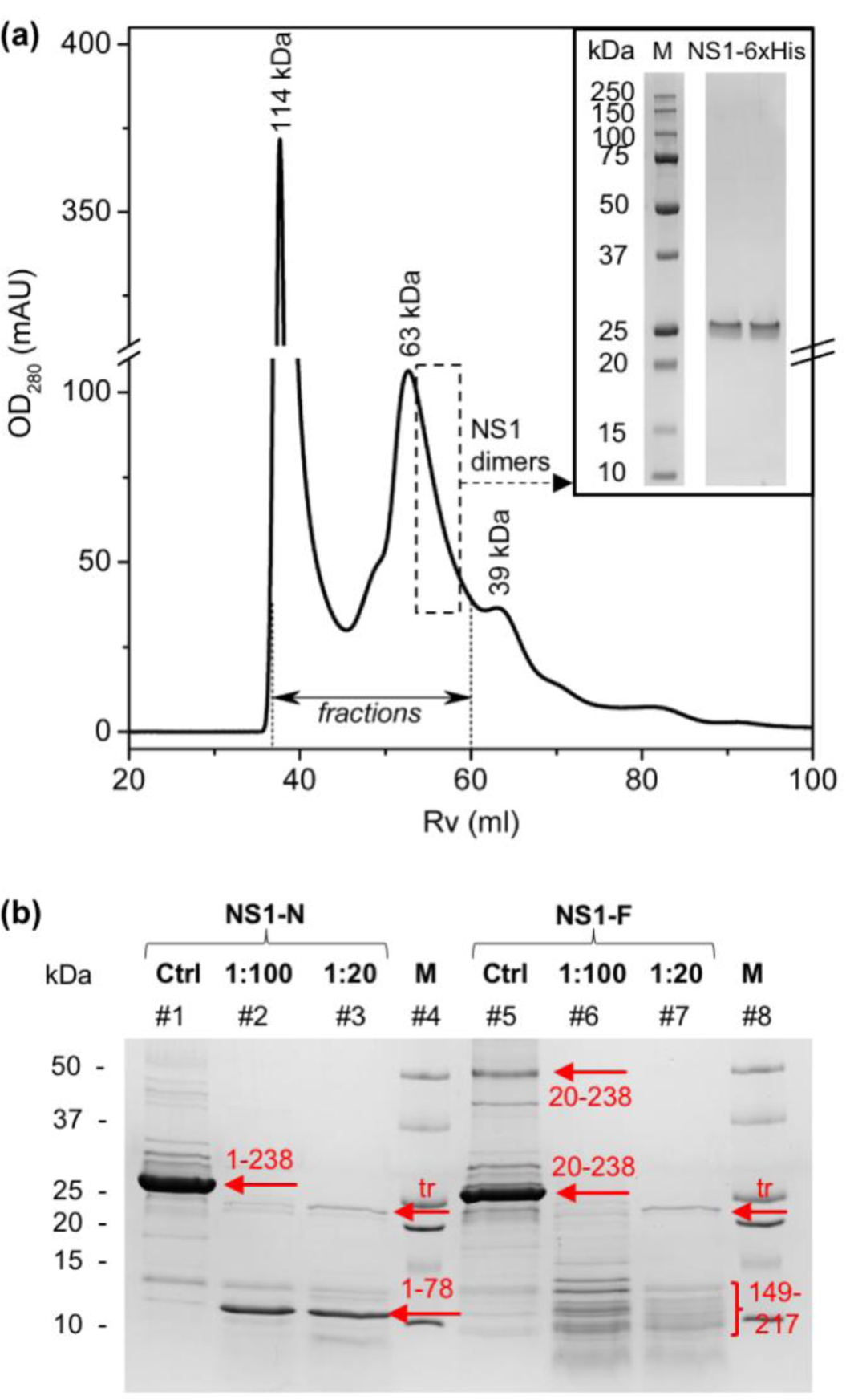
Protein isolation and size analysis. **(a)** – Gel-filtration results. Inset: SDS-PAGE of purified recombinant NS1-6xHis protein. Using a calibration curve, approximate molecular masses corresponding to each peak were identified. Rv – retention volume, OD_280_ – optical density at 280 nm, M – molecular mass standard. **(b)** – SDS-PAGE of NS1-N and NS1-F after protease treatment at two protease:protein ratios (1:100 and 1:20). Ctrl – NS1-N or NS1-F incubated without trypsin, under the same conditions. Arrows and numbers indicate bands and their mass spectrometry-identified NS1-6xHis a.a. residues. The band corresponding to trypsin is marked (tr).

It was shown that it is possible to form amyloid-like fibrils by incubating the isolated protein (at a concentration of 3 mg/ml in 2x phosphate-buffered saline) for 66 hours at a temperature of 55°C. The presence of protein aggregates was shown using electron microscopy (Figure 2).

**Figure 2.**
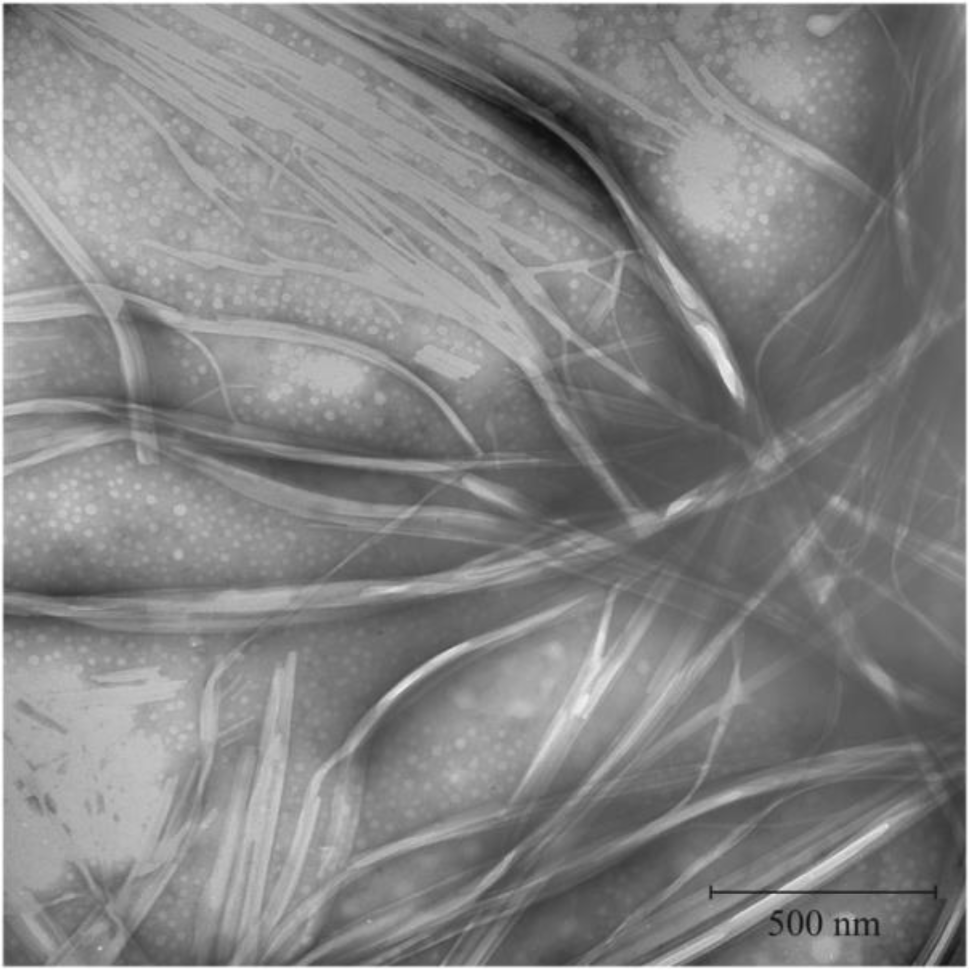
Electron micrograph showing recombinant protein aggregates. The scale bar is 500 nm

The ability of the recombinant NS1 protein to bind amyloid-specific dyes (Congo red and Thioflavin T) supports the conclusion that the detected fibrillar aggregates are amyloid in nature (Figure 3). The fluorescence quantum yield of NS1-F with ThT exceeds that of pure ThT by 12-fold (Figure 3a). In addition, in the CR/NS1-F ‘difference spectrum’, a 540 nm maximum is seen (Figure 3b), which indicates NS1-F protein complexes binds CR, which is characteristic of amyloid-like fibrils. It should be noted that interaction of NS1-N with ThT or CR was not observed (data not shown).

**Figure 3.**
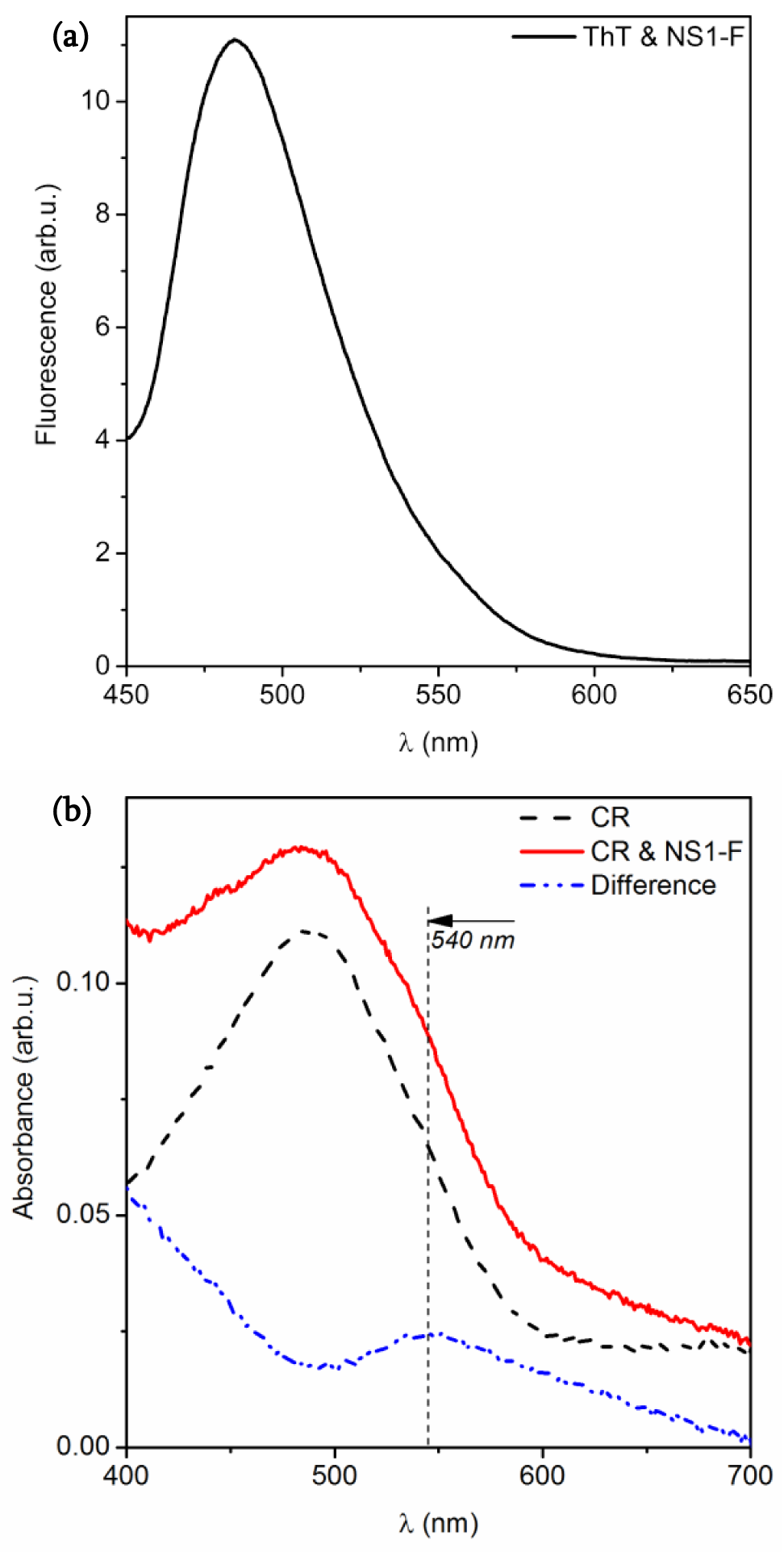
Spectra of NS1 fibrillar aggregate complexes. NS1-F with **(a)** ThT (fluorescence) and with **(b)** CR (absorbance).

Using proteolysis followed by SDS-PAGE and mass spectrometry, it was shown that amino acid sequence accessibility of native (NS1-N) and fibrillar (NS1-F) protein forms to trypsin differed. Native and fibrillar NS1-6xHis protein forms were subjected for tryptic hydrolysis for 24 hours at 37°C. Mock-treated and cleaved protein samples were analyzed by SDS-PAGE followed by mass spectrometry (Fig.1b). Protease treatment of NS1-N, at both trypsin concentrations (Fig.1b, lanes 2,3), resulted in one major protein band (11 kDa) containing an NS1-6xHis fragment representing amino acid residues (a.a.r.) 1 to 78 according to mass spectrometry data (Score 116/70). It should be noted that additional weak bands appear in control samples (for both NS1-N and NS1-F). These likely arise during the incubation for the preparations: 37°C for NS1-N; and 55°C followed by 37°C for NS1-F. Therefore, we limit our analysis and conclusions to only those bands which substantially differ (presence or intensity) from those present in controls. It can be seen in Fig.1b (lane 5) that a band with electrophoretic mobility corresponding to 54 kDa appeared after NS1 fibril formation. NS1-6xHis was reliable identified in that band (Score 274/70). It can be concluded that the NS1-F sample contains SDS-resistant NS1-6xHis dimers. The presence of such protein forms is characteristic of amyloid-like fibrils.

In the case of NS1-F trypsin hydrolysis, the resultant protein fragments possessed a wide range of molecular masses (Fig.1b, lane 6). At the higher trypsin concentration (lane 7), the set of fragments became more limited, generally in the 10-13 kDa range. In this molecular mass area, the NS1-6xHis fragment from a.a.r. 149 to 217 was preliminary identified (Score 76/55). It should be noted that small amounts of bigger protein fragments were also found in that PAGE area, as identified by mass spectrometry: a.a.r. 118-217; and a.a.r. 118-217 with 22-78. Together, these identified fragments cover almost the entire protein sequence. This indicates that NS1-F is hydrolyzed by trypsin without any specific resistant regions.

Thus, it can be concluded that the N-terminal, native-form NS1-6xHis fragment (a.a.r. 1-78) remains intact following trypsin digestion (Table 1, green). In the fibrillar NS1-6xHis form, electrophoresis does not indicate a major stable region like in the native form. Multiple bands of lesser intensity occur. These were identified, however, as being C-terminal fragments featuring slight trypsin resistance, corresponding to a.a.r. 149-217 (Table 1, red). It is interesting that the fibrillar NS1 form was found to be more protease susceptible, and therefore accessible, than the native form.

It should be noted that, when packaged as a crystalline structure, the indicated protease-resistant portion (a.a.r. 1-78) is located in the monomer-monomer contact area indicated by analysis of the PDB file [21] (Figure 4). The trypsin accessibility of the NS1 region in fibrillar form, in contrast to native form, demonstrates that significant conformational transitions occur during fibril formation, leading to distortion of native functional dimers. In addition, the a.a.r. 149-217 region is located near the protein’s beta-domain (Figure 4, inset) and partially overlaps it, which may indicate a potential role of the beta-domain in the NS1 fibril-formation process.

**Figure 4.**
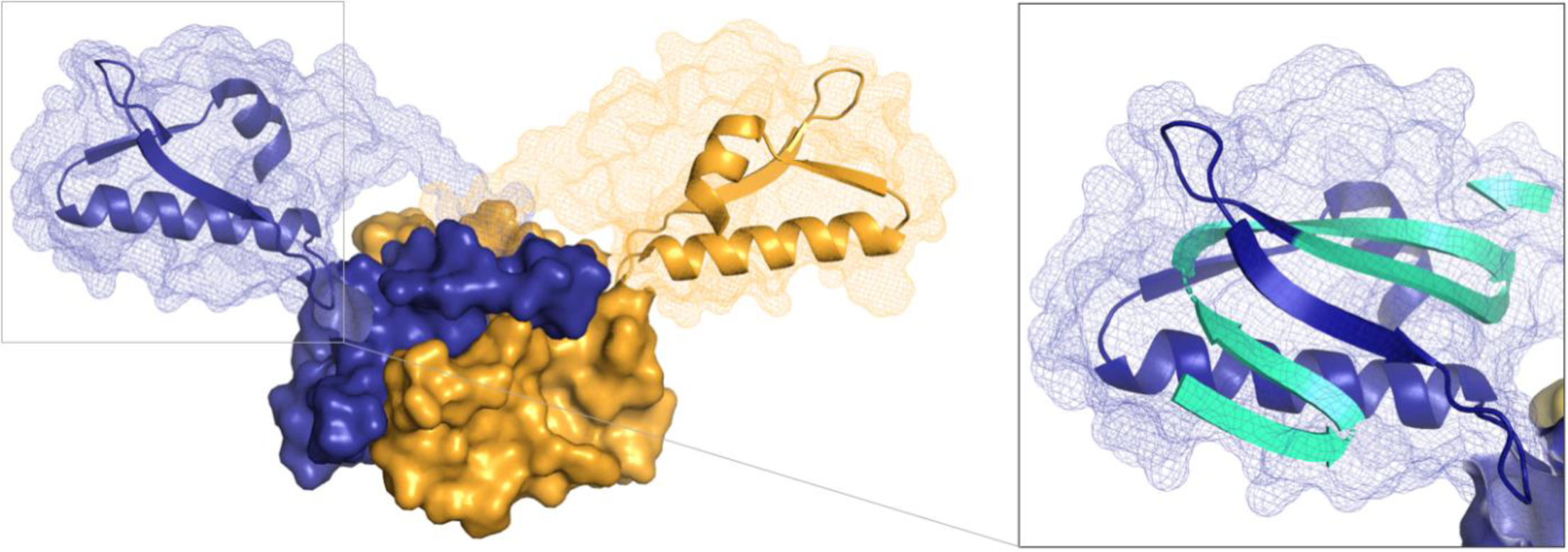
Structural model of NS1 monomer interaction obtained by PDB (4OPH) data analysis. Monomers are indicated in dark blue and yellow. Areas homologous to NS1-N a.a.r. 1-78 (resistant to protease treatment) and to NS1-F a.a.r.149-217 are shown using surface and cartoon representation, respectively. Inset: localization of a.a.r. 149-217 (dark blue, cartoon representation) near NS1’s beta-domain (cyan and dark blue arrows, cartoon representation).

## CONCLUSIONS

The data obtained, together with published data on influenza A NS1 protein’s tendency towards structural polymorphism, indicate that NS1 can form amyloid-like fibrils *in vitro*. Further work will be needed to see if these structures are possible or relevant *in vivo*. The formation of such fibrils may play a role in viral infection by recruiting cellular factors featuring sequence homology with NS1, for example by a prion-like mechanism. Experiments and screening for identification of amyloid-like protein aggregates in influenza-infected cells are planned. Recombinant NS1 protein has the ability to form amyloid-like fibrils. During fibril formation, SDS-resistant dimers appear.

The conformation of such dimers differs from that of native, functional NS1 protein dimers. This fact is confirmed by significant differences in the amino acid residues exposed and susceptible to protease hydrolysis. Presumably, the NS1 beta domain plays an important role in non-native dimers and, hence, fibrillogenesis. In infected cells, NS1 fibrillogenesis may play a role in signaling by prion-like mechanisms or in recruitment of cellular antiviral factors.

## ACKNOWLEDGMENT

This work was supported by the NRC “Kurchatov Institute” (N° 1363).

